# PhISCS - A Combinatorial Approach for Sub-perfect Tumor Phylogeny Reconstruction via Integrative use of Single Cell and Bulk Sequencing Data

**DOI:** 10.1101/376996

**Authors:** Salem Malikic, Simone Ciccolella, Farid Rashidi Mehrabadi, Camir Ricketts, Khaledur Rahman, Ehsan Haghshenas, Daniel Seidman, Faraz Hach, Iman Hajirasouliha, S. Cenk Sahinalp

## Abstract

Recent technological advances in single cell sequencing (SCS) provide high resolution data for studying intra-tumor heterogeneity and tumor evolution. Available computational methods for tumor phylogeny inference via SCS typically aim to identify the most likely *perfect phylogeny tree* satisfying *infinite sites assumption* (ISA). However limitations of SCS technologies such as frequent allele dropout or highly variable sequence coverage, commonly result in mutational call errors and prohibit a perfect phylogeny. In addition, ISA violations are commonly observed in tumor phylogenies due to the loss of heterozygosity, deletions and convergent evolution. In order to address such limitations, we, for the first time, introduce a new combinatorial formulation that integrates single cell sequencing data with matching bulk sequencing data, with the objective of minimizing a linear combination of (i) potential false negatives (due to e.g. allele dropout or variance in sequence coverage) and (ii) potential false positives (due to e.g. read errors) among mutation calls, as well as (iii) the number of mutations that violate ISA - to define the *optimal sub-perfect phylogeny.* Our formulation ensures that several lineage constraints imposed by the use of variant allele frequencies (VAFs, derived from bulk sequence data) are satisfied. We express our formulation both in the form of an integer linear program (ILP) and - for the first time in the context of tumor phylogeny reconstruction - a boolean constraint satisfaction problem (CSP) and solve them by leveraging state-of-the-art ILP/CSP solvers. The resulting method, which we name PhISCS, is the first to integrate SCS and bulk sequencing data under the finite sites model. Using several simulated and real SCS data sets, we demonstrate that PhISCS is not only more general but also more accurate than the alternative tumor phylogeny inference tools. PhISCS is very fast especially when its CSP based variant is used returns the optimal solution, except in rare instances for which it provides an optimality gap. PhISCS is available at https://github.com/haghshenas/PhISCS.

## 1 Introduction

The clonal theory of cancer evolution suggests that cancer is an evolutionary disease where multiple distinct cellular populations (i.e. subclones) emerge through successive rounds of mutation and selection. At the time of clinical diagnosis, most tumors are heterogeneous, consisting of multiple subclones harboring different sets of somatic mutations. An increasing evidence suggests that this phenomenon, better known as “intra-tumor heterogeneity” (ITH), has a profound impact on treatment outcomes and that the existence of treatment resistant subclones is one of the main causes of treatment failures [1]. Deciphering intra-tumor heterogeneity and tumor evolutionary history thus represent some of the key challenges in designing efficiently combined therapies and better understanding of dynamics of cancer initiation and progression.

Most of the existing approaches for studying ITH are based on analyzing data from next-generation bulk sequencing experiments where only an average signal over a large number of cells is obtained. In the past few years, numerous computational methods for analyzing such signals with the aim of inferring tumor subclonal composition and evolutionary history have been developed [2, 3, 4, 5, 6, 7, 8, 9, 10, 11, 12, 13]. Even though these methods employ a variety of computational approaches – each with a particular strength, all have theoretical limitations, mainly due to the limited resolution offered by bulk sequencing data.

While it is still expensive and experimentally challenging to robustly perform single cell library preparation, recent technological advancements in single-cell sequencing (SCS) potentially provide higher resolution data for studying ITH. Single-cell datasets are, however, characterized by high levels of sequencing noise that includes both false positive (e.g. due to the read errors) and false negative (e.g. due to the allele dropout or variance in sequence coverage) mutation calls, as well as *missing* values for mutations from sites affected by DNA amplification failure.^1^ This necessitates the development of sophisticated computational methods that are sensitive to the noise characteristics of SCS data, while incorporating the assumptions of the clonal theory of cancer evolution to tumor evolution modeling.

Available methods for studying ITH via the use of SCS data are all based on probabilistic approaches with the goal of inferring the *most-likely perfect-phylogeny* for a tumor. SCITE [15], for example, is a Markov Chain Monte Carlo search method that aims to infer the maximum-likelihood (ML) mutational history from a potentially incomplete and noisy matrix containing genotypes of single cells. OncoNem [16] is a maximum likelihood based search approach to identify homogeneous cellular subpopulations and infer both their genotypes and the tree describing their evolutionary history. For achieving their respective goals SCITE and OncoNem both rely on the *infinite sites assumption* (ISA), i.e. that each genomic position is affected by at most one mutation hit in the entire tumor phylogeny. A more recent maximum likelihood based approach, SiFit [17], aims to extend the above by employing a finite sites model of evolution that accounts for deletions, loss of heterozygosity (LOH) and point mutations on genomic sites. However, none of the above approaches provide means to integratively use SCS with bulk sequencing data, which, in principle may provide additional guidance to tumor phylogeny construction process. Another recent tool, ddClone [18] is the first to combine the strengths of bulk and SCS data in a joint statistical inference model for the most likely tumor subclonal composition. However, ddClone does not aim to build a tumor phylogeny and is not suitable to study cancer evolution. Finally, some of us developed B-SCITE [19] with the aim of integrating SCITE with CITUP [9] so as to make joint use of SCS and bulk sequencing data. B-SCITE is an MCMC based tool, and as per SCITE it does not account for ISA violations.

Even though the above methods for SCS data analysis are probabilistic, many of the related methods for bulk sequencing data analysis are combinatorial in nature [2, 4, 7, 8, 9, 11]. Combinatorial, in particular integer linear programming (ILP), formulations for phylogeny inference can be traced back to *Haplotype Inference Problem* (HIP) [20]. Given a binary incomplete matrix M of n rows (corresponding to *species)* and m columns (corresponding to *sites),* HIP asks to decompose each row to two binary vectors (haplotypes) so that the haplotypes can fit in a *Perfect Phylogeny,* i. e. a phylogeny satisfying ISA. This problem can be formulated and efficiently solved as an instance of ILP. Later, a similar formulation was proposed in [21] to solve the *Persistent Phylogeny Problem* [22, 23]. In this problem, the goal is to compute a persistent phylogeny that is defined as a phylogeny in which each mutation is allowed to be “lost” at most once. Recently, an extension of formulation from [21] was proposed in [24], where more general phylogeny models are used and the goal is to infer entire cancer phylogenies by the use of bulk sequencing data.

ILP formulations for HIP and related problems are routinely solved through commercial tools such as Gurobi or IBM CPLEX - which have been developed over many years and provide reliable and fast solutions for relatively small sized optimization problems. These solvers aim to optimize a typically linear objective while satisfying a number of numerical constraints. As such, ILP is related to another fundamental problem, the *boolean Constraint Satisfaction Problem* (CSP) that can be used as an alternative for modeling many ILP problems encountered in practice.

Perhaps the best-known variant of CSP is the satisfiability problem (SAT) which asks to find a boolean assignment to a set of input variables to satisfy (the conjunction of) a number of boolean constraints.^2^ Among other variants, Max-SAT asks to find a boolean assignment to variables such that not necessarily all but the maximum number of input constraints are satisfied, while weighted version of Max-SAT, which can be abbreviated as wMax-SAT, asks for the assignment that maximizes the sum of (user defined) weights of the constraints satisfied. The generality of SAT and (w)Max-SAT has prompted the development of many tools to solve them with the goal of obtaining solutions to practical instances of NP-complete problems. These tools compete in the annual SAT conference on several benchmarking datasets generated by a wide variety of applications (see http://sat2017.gitlab.io). Recently developed wMax-SAT solvers such as Maxino [25] and MaxHS [26, 27, 28], are not only very fast but the later is also open source. As a result, a number of studies demonstrated the utility of CSP solvers for the haplotype inference problem and its variants - before the advent of high throughput sequencing [29, 30, 31]. To the best of our knowledge, however, no study has explored the use of CSP in the context of intra-tumor heterogeneity or tumor phylogeny modeling.

### Our Contributions

In this paper, we introduce three novel combinatorial formulations for inferring tumor phylogenies via an integrative use of single-cell and bulk sequencing data. (1) Our simplest formulation asks to minimize a weighted sum of potential false negative (which are common) and false positive (which are rare) mutation calls in genotypes of single cells, whose correction will result in a perfect phylogeny. (2) The goal of our more general formulation is to compute a *sub-perfect* phylogeny, which not only requires such mutation calls to be corrected but also needs the elimination of (at most a user defined number of) mutations that violate ISA (e.g. due to LOH - and are relatively rare). More specifically, this formulation asks to minimize a weighted sum of mutations to be corrected, given an upper bound on the number of mutations to be eliminated (due to ISA violations) in order to achieve a perfect phylogeny. (3) Our most sophisticated formulation has additional constraints imposed by the use of variant allele frequencies (VAFs) of single nucleotide variants (from regions not affected by copy number aberrations) that can be estimated from bulk sequencing data (as a proxy to the cellular prevalence of a given mutation). These *lineage constraints* impose ancestor-descendant dependencies among mutation pairs (e.g. the prevalence of an ancestral mutation cannot be lower than that of a descendant) or triplets (e.g. the prevalence of an ancestral mutation cannot be lower than the sum of two descendant siblings) and improve inference accuracy. We describe solutions to each of the three formulations to address problems of varying complexity and data availability (i.e. some data sets have no ISA violations and some do not come with matching bulk sequencing data).

We name our general formulation and the resulting program PhISCS (Ph*ylogeny of tumors using Integrated bulk and Single Cell Sequencing data),* which comes in two flavors: (i) PhISCS-1 expresses our formulation in the form of an ILP and efficiently solves it by the use of Gurobi Optimizer. (ii) PhISCS-B expresses our formulation in the form of a Boolean CSP and solves it by the use of open source solvers for wMax-SAT such as MaxHS - many times more efficiently than PhISCS-I.

Many of the available tools for studying intra-tumor heterogeneity formulate the problem as an ILP or quadratic integer programming (QIP) and solve it via commercial tools such as Gurobi or CPLEX. Our CSP formulation (specifically in wMax-SAT) is the first to express a tumor phylogeny reconstruction problem combinatorially, but in a form other than ILP/QIP. Additionally, unlike many alternatives, PhISCS has the ability to integrate single cell and bulk sequencing data, and can simultaneously infer tumor phylogeny and clonal composition of the tumor sample. Furthermore, recent studies suggest that ISA, that forms the basis for most of the above tools (with SiFiT being the main exception), could be violated to some degree in tumor phylogenies [17, 32] making it impossible to establish a perfect phylogeny. PhISCS addresses this issue by eliminating (a small number of) mutations that violate ISA (with a cost reflected in the objective) and solves the tumor phylogeny reconstruction problem for both simulated and real data, more efficiently and more accurately (clearly for simulations but also real data) than the available alternatives.

Our final contribution is on assessing the (dis)similarity between two tumor phylogenies - typically between G, the ground truth tree, and T, the tree inferred by any method. Commonly used measures of similarity between tumor phylogenies such as *lineage consistency*, and *non-lineage consistency* (used by [9] and others), are defined based on the proportion of mutation pairs with the same lineage relationship in the two trees fail to capture fundamental topological differences between simulated ground truth and inferred trees, especially of different levels of *granularity* [33]. The more recent *co-clustering consistency*, defined as a function of the proportion of mutation pairs that are differentially clustered in the two trees, suffers from the same problem [33]. In order to overcome the limitations of the available measures, some of us very recently introduced *Multi-labeled tree edit distance* (MLTED), the minimum number of label deletions, accompanied with (an arbitrary number of) empty leaf deletions and vertex expansions, to transform each of two trees to a maximal sized common tree. MLTED successfully captures the differences between tumor phylogenies of any granularity. In this paper, we also introduce *Heuristic-based Multi-labeled tree edit distance* (HMLTED) which, as demonstrated below, accurately captures the dissimilarity between a ground truth tree and an inferred tree in all our simulations.

## 2 Methods

In this section, we formulate our integrative tumor phylogeny reconstruction as a combinatorial optimization problem. We first discuss two special cases of the problem for the case when only single cell sequencing data is available. i.e. (i) a special case where the ISA cannot be violated (Section 2.1) (ii) the case where ISA can be violated (Section 2.2). We then describe the general integrative problem where both bulk and SCS data are available. We present solutions for this problem using a novel Integer Linear Program (ILP) as well as a Constraint Satisfaction Program (CSP).

### 2.1 Only single cell data and no ISA violation

The input is a ternary matrix *I* with *n* rows and *m* columns, where columns represent mutations and rows represent genotypes of single cells observed in a single-cell sequencing experiment. The entry *I*(*i,j*) = 0 indicates the absence, *I*(*i,j*) = 1 indicates the presence and *I*(*i,j*) = 2 indicates the lack of knowledge about absence or presence (i.e. missing entry) of a mutation *j* in a cell *i*.

We ask to find the minimum weighted number of *bit flips* (typically from 0 to 1 and rarely from 1 to 0) and *bit assignments* (assigning a 0 or 1 to an entry with value 2), where *bit assignments* are not a part of the objective, such that the resulting matrix provides a Perfect Phylogeny (PP). We recall that a binary matrix is a PP if the *three-gametes rule* holds, i.e. for any given pair of columns (mutations) there are no three rows (cells) with configuration (1, 0), (0,1) and (1,1). *Bit flipping* in the input matrix *I* is essential to building a PP as some mutation inferences in I are false positives and some mutations are not indicated in *I* (false negatives) as they do not have sufficient read support in sequenced single cells.

To allow correction of noisy genotypes in *I* (i.e. bit flips and bit assignments), for each cell *i* and mutation *j*, we introduce a binary variable *Y*(*i,j*) which denotes the (unknown) true status (i.e. absence or presence) of the mutation *j* in the cell *i*. Assuming that *α* and *β* respectively denote false positive and false negative error rates of single-cell data we use the following scoring scheme:

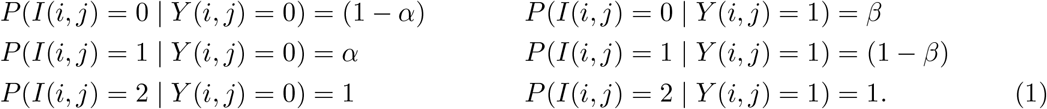

The above scoring scheme can be rewritten as:

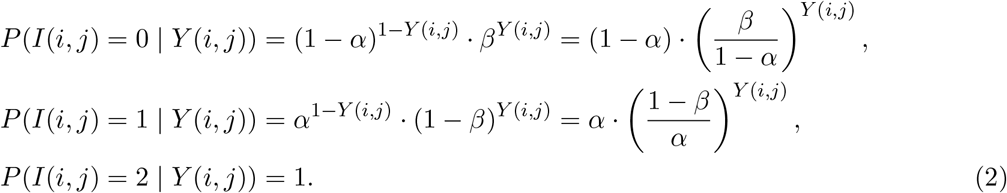

The likelihood of an arbitrary conflict free matrix *Y* is defined as:

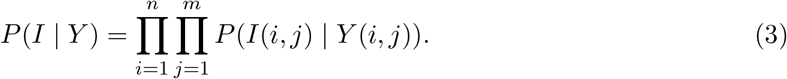

Here our goal is to find a conflict free matrix *Y* such that the likelihood defined in 3 is maximized. This is equivalent to maximizing logarithm of *P*(*I | Y*) which is equal to:

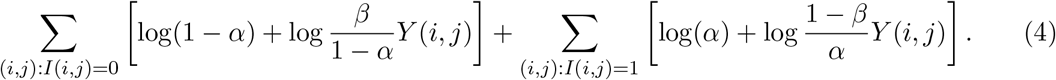

In order to enforce that matrix *Y* satisfies the three-gametes rule, for each pair of mutations (*p, q*), we first introduce variables *B*(*p, q, a, b*), for each (*a, b*) ∈ {(0,1), (1,0), (1,1)}. The variable *B*(*p,q,a,b*) is set to 1 if and only if there exists row *r* such that *Y*(*r,p*) = *a* and *Y*(*r,q*) = *b*. This property of matrix B is guaranteed by adding the following constraints for all 1 ≤ *i* ≤ *n* and 1 ≤ *p*, *q* ≤ *m*:

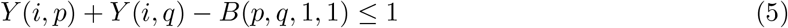

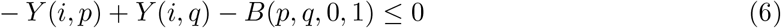

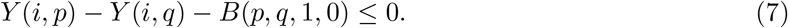

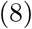

Now, adding constraints

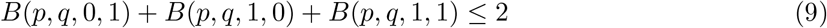

for all 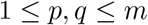 suffices to ensure that three-gametes rule holds for matrix *Y*.

The problem defined above represents an instance of ILP and can be solved using any of the standard ILP solvers.

### 2.2 Allowing for ISA violations in the model

As we have already discussed in Section 1, recent evidence suggests ISA might be violated for a subset of mutations in the input data. To account for this, we introduce a more general version of what we discussed in the previous section where we allow *elimination* (i.e. deletion) from the input matrix of a given maximum number of mutations which do not satisfy ISA, while the remaining mutations, after genotype corrections, are expected to satisfy PP. In order to achieve this, for each mutation *q* we introduce binary variable *K*(*q*) which is set to 1 if and only if mutation *q* is among eliminated mutations. To exclude eliminated mutations from three-gametes rule, we do not require mutational pairs (*p,q*), where at least one of *p* and *q* is among eliminated mutations, to fulfill this rule. Therefore we modify constraint 9 from the integer linear program we described in Section 2.1 by replacing it with:

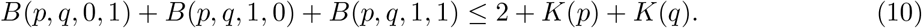

The objective defined in 4 is also modified so that the eliminated mutations don’t contribute to the objective score. This leads to the following objective for the case allowing for ISA violations:

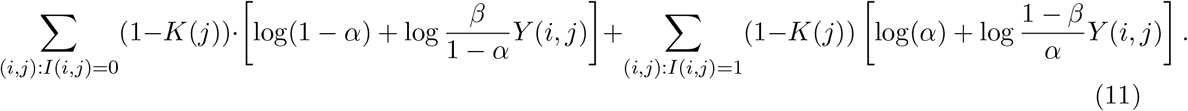

All other constraints used previously in the limited version of the problem are preserved. The latest objective contains quadratic terms of the form *K*(*j*)*Y*(*i,j*) which can be transformed to the linear variables using standard linearization techniques. One can observe that mutation elimination never decreases data likelihood hence the global optimum in the above maximization problem is achieved when all variables *K* are set to 1. However, in real applications we expect only a limited number of ISA violating mutations and therefore set the upper bound *k_max_* on the number of eliminated mutations which is implemented by the addition of the following constraint

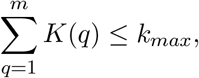

where *k_max_* is empirically determined constant.

### 2.6 Additional ILP Constraints to Integrate VAFs Derived from Bulk Sequencing Data with SCS

We can integrate SCS data with bulk sequencing data - specifically VAF of each mutation we consider - through additional linear constraints. These constraints will only apply to the set of single nucleotide variants from the regions not affected by copy number aberrations. Suppose that a particular SNV, denoted *M*, satisfies the above requirement; let *υ* and *r* respectively denote the number of reads supporting the variant and the reference allele at the genomic locus of *M*. The VAF of *M* is typically defined as 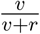. Since below we are interested in *cellular prevalence* rather than the VAF below, we define 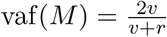. (Cellular prevalence represents the expected fraction of cells in the sample that harbor *M*.)

Before defining constraints related to VAFs, we first define the *root node* via a new row, indexed by 0, that represents genotype of a healthy cell. We also add a new column, indexed by 0, and associated *null mutation M*_0_ which represents mutation specific to the normal cell or, in other words, germline SNP present in all cells. We set I(t, 0) = 1 for *t* = 0,1…,*n* and *I*(0,*p*) = 0 for *p* = 1,2,…, *m*. We also set vaf(*M*_0_) = 1 and do not allow elimination of *M*_0_. Matrices *B* and *Y* are also expanded in an obvious way by allowing mutational indices to be equal to 0. The remainder of the tree topology is imposed through additional constraints that specify ancestor-descendent relationships in a consistent manner across all nodes:

1. We must satisfy the following constraints that are trivially converted into boolean expressions: (i) *K* (0) = 0, (ii) *Y* (*t*, 0) = 1 for *t* = 0, 1,…,*n*, and (iii) *Y* (0,*p*) = 0 for *p* = 1, 2,…,*m*.
2. If a mutation *p* is an ancestor of a mutation *q* and ISA holds for both *p* and *q* then the true cellular prevalence of *p* is expected to be greater than or equal to true cellular prevalence of *q*. Since vaf(*p*) and vaf(*q*) reflect cellular prevalences as discussed above, we expect that in the implied evolutionary tree vaf(*p*)(1 + *δ*) ≥ vaf(*q*), where *δ* is some positive constant which allows for the noise typically present in the observed VAFs. In order to incorporate VAFs in our model, we introduce binary function *α*, such that *α*(*p, q*) = 1 only if *p* is an “ancestor” of *q*. By definition we set *a*(*p,p*) = 0 for all *p* ∈ {0,1,…, *m*}. The constraints that we need to introduce are thus as follows.

(a) For all pairs of mutations p, q we must satisfy:

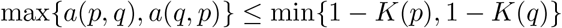

here, for any mutation *r, K(r)* = 1 indicates that the column *r* in input matrix *I* has been eliminated.
(b) Each non-eliminated mutation *q* different from null mutation must have at least one ancestor. This is ensured by adding the following constraint:

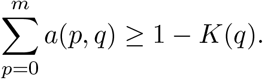 On the other hand, null mutation has no ancestors so we set *a*(*p*, 0) = 0 for all *p* ∈ {0, 1, . . . , *m*}.
(c) Consider two non-eliminated mutations *p* and *q*. If *a*(*p, q*) = 1 then in genotype corrected output matrix *Y* the column *p* should dominate the column *q* - i.e. for each cell/row *r* if the entry for *p* is 0 then the entry for *q* should also be 0. In other words, there should not exist row *r* such that *Y*(*r,p*) = 0 and *Y*(*r, q*) = 1, which is equivalent to *B*(*p, q*, 0, 1) = 0. To ensure this, for all pairs of mutation (*p, q*), we add the following constraint:

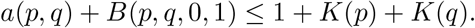
(d) If, for two non-eliminated mutations *p* and *q*, matrix *Y* contains cell in which *p* is present and *q* is absent (i.e. there exists *i* such that *Y*(*i,p*) = 1 and *Y*(*i, q*) = 0, which is equivalent to *B*(*p, q,* 1,0) = 1), as well as cell where both *p* and *q* are present (i.e. there exists *j* such that *Y*(*j,p*) = 1 and *Y*(*j, q*) = 1, which is equivalent to *B*(*p,q,* 1,1) = 1), then p must be ancestor of *q* (i.e. *a*(*p,q*) = 1). In order to ensure this, for all pairs of mutations (*p, q*) we add the following constraints:

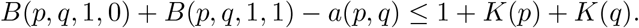
(e) For some small user defined error tolerance value *δ* > 0 that accounts for variation in sequence coverage, if vaf(*q*) > vaf(*p*)(1 + *δ*) then *a*(*p,q*) = 0; in other words for every pair of mutations *p* and *q* we must satisfy:

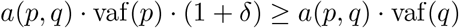
(f) For all triplet of mutations *p, q, r*, we must ensure that if *a*(*p, q*) = 1 and *a*(*q, r*) = 1 then *a*(*p, r*) = 1:

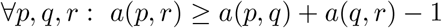
3. For all triplet of mutations *p, q* and *r* (such that *p* is an ancestor of *q* and *r* but *q* and *r* do not have an ancestor descendant relationship, i.e. *a*(*p, q*) = *a*(*p, r*) = 1 and *a*(*q, r*) = *a*(*r, q*) = 0) we must satisfy:

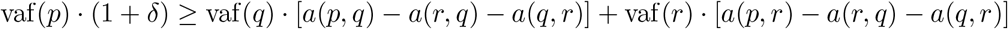

### 2.4 PhISCS-B for Tumor Phylogeny Inference via SCS

In this section we show how to reduce the ILP provided in Section 2.1 to a wMax-SAT problem. For each input entry *I*(*i,j*), 1 ≤ *i* ≤ *n*, 1 ≤ *j* ≤ *m*, our goal is to compute the value *X*(*i,j*), which indicates whether *I* (*i,j*) is flipped or not (we will also initially set all entries *I* (*i,j*) = 2 to 0, but will have no penalty for flipping their values to 1 later) resulting in the output matrix *Y*, where *Y* (*i,j*) = *X* (*i,j*) ⊕ *I* (*i,j*), which admits a PP. In order to achieve this we use a set of additional variables *B*(*p, q, a, b*) (see Section 2.1) that need to satisfy the following *hard* constraints (the constraints that *need* to be satisfied):

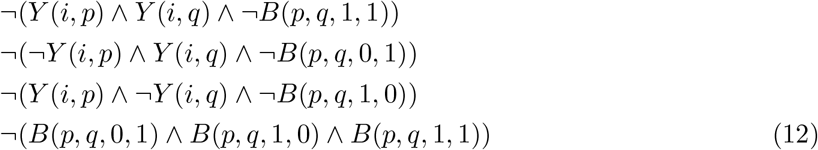

We can now define our objective as satisfying all the hard constraints with alterations on the input matrix *I* with maximum probability, where each alteration (indicating a false positive or false negative) is independent. This objective corresponds to the minimizing the (weighted) number of flipped entries in the solution matrix *Y* in comparison to *I*, or, for the purpose of formulating the problem as an instance of wMax-SAT, maximizing the weighted sum of the following “soft” constraints (for all *i,j*, s.t. *I* (*i,j*) ≠ 2 originally):

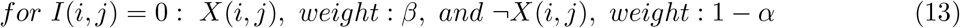

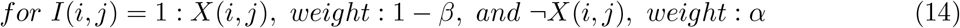

In order to account for ISA violations, for each column *j* ∈ {1,2,…, *m*} we introduce boolean variable *K*(*j*) that is set to 1 if and only if column *j* is eliminated (i.e. mutation corresponding to column *j* is not considered as a part of the output).

Similarly as in 2.2, we allow at most *k_max_* columns to be eliminated, where *k_max_* is a user-defined constant. In order to ensure that no more than *k_max_* of variables *K*(1), *K*(2),…, *K*(*m*) are set to 1, for each possible (*k_max_* + 1)-tuple (*i*_1_, *i*_2_,…, *i_k_max+1__*) of integers such that 1 ≤ *i*_1_ < *i*_2_ < … < *i_k_max_+1_* ≤ *n* we add the the following hard clause

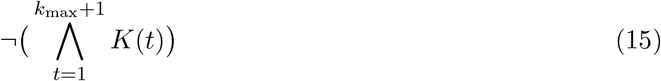

to our model. Now, for any eliminated column *p* we do not have to check whether it is in conflict with any other column *q* or vice versa. Therefore, for each pair (*p, q*) of columns we replace the constraint 12 above with the following.

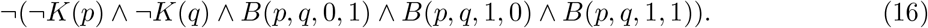

### 2.5 Additional Boolean Constraints to Integrate Bulk Sequencing with Single Cell Seq Data

In order to integrate information derived from bulk sequencing data, represented in the form of VAFs of the given set of mutations (see Section 2.3 for details), we explicitly impose a tree structure on the output matrix *Y* through the use of a number of boolean constraints.

The boolean constraints below start by defining the *root node* via a new row, indexed by 0, that represents genotype of a healthy cell. We also add a new column, indexed by 0, and associated *null mutation M*_0_ which represents mutation specific to the normal cell or, in other words, germline SNP present in all cells. We set *I*(*t*, 0) = 1 for *t* = 0,1…, *n* and *I*(0,*p*) = 0 for *p* = 1,2,…, *m*. We also set vaf(*M*_0_) = 1 and do not allow elimination of *M*_0_. The remainder of the tree topology is imposed through additional constraints that specify ancestor-descendent relationships in a consistent manner across all nodes:

1. We must satisfy the following constraints that are trivially converted into boolean expressions: (i) *K* (0) = 0, (ii) *Y* (*t*, 0) = 1 for *t* = 0,1,…,*n*, and (iii) *Y* (0,*p*) = 0 for *p* = 1, 2,…,*m*.
2. If a mutation *p* is an ancestor of mutation *q* in the implied evolutionary tree, then vaf(*p*) ≥ vaf(*q*) must be satisfied, within some error tolerence that can usually be assumed to be small for the existing datasets containing both, single cell and bulk sequencing data. Namely, in such datasets, bulk sequencing data is typically of deep coverage resulting in higly accurate and reliable estimates of mutational VAFs. In order to exploit values of VAFs using the above dependency between phylogenetic relation of mutational pairs and their VAFs, we introduce boolean function a, such that *a*(*p, q*) = 1 if and only if *p* is an “ancestor” of *q*. The hard constraints that need to be imposed on a are as follows.

(a) For all pairs of mutations *p* and *q*, where both *p* and *q* are different from null mutation, we must satisfy:

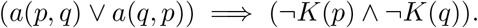
(b) For all non-eliminated mutations *q* different from null mutation we must make sure that it has an ancestor mutation (which could be null mutation). This is achieved by imposing the following constraint:

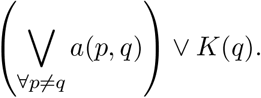
(c) Consider two non-eliminated mutations *p* and *q*. If *a*(*p, q*) = 1 then in genotype corrected output matrix *Y* the column *p* should dominate the column *q* - i.e. for each cell/row *r* if the entry for *p* is 0 then the entry for *q* should also be 0. In other words, there should not exist row *r* such that *Y*(*r,p*) = 0 and *Y*(*r, q*) = 1, which is equivalent to *B*(*p, q*, 0,1) = 0. To ensure this, for all pairs of mutation (*p, q*), we add the following constraint:

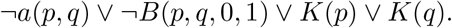
(d) If, for two non-eliminated mutations *p* and *q*, matrix *Y* contains cell in which *p* is present and *q* is absent (i.e. there exists *i* such that *Y*(*i,p*) = 1 and *Y*(*i, q*) = 0, which is equivalent to *B*(*p, q*, 1,0) = 1), as well as cell where both *p* and *q* are present (i.e. there exists *j* such that *Y*(*j,p*) = 1 and *Y*(*j, q*) = 1, which is equivalent to *B*(*p, q*, 1,1) = 1), then *p* must be ancestor of *q* (i.e. α(*p, q*) = 1). In order to ensure this, for all pairs of mutations (*p, q*) we add the following constraints:

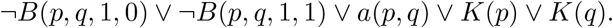
(e) For some small user defined error tolerance value *δ* > 0 that accounts for variation in bulk sequencing coverage, if vaf(*q*) > vaf(*p*) · (1 + *δ*) then *a*(*p, q*) = 0; in other words for each pair of mutations *p* and *q* for which *a*(*p, q*) = 1, we must satisfy vaf(*p*) · (1 + *δ*) ≥ vaf(*q*). In order to express this as a boolean constraint we introduce a new boolean function Pvaf(*p, q*) defined for all pairs of mutations *p* and *q* (as a part of the input specification) as follows:

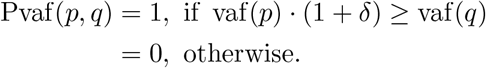 Then the constraint that must be satisfied for each pair of mutations *p* and *q* is:

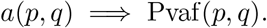
(f) For all pairs of mutations *p* and *q* (including null mutation) we must ensure that, if *p* dominates *q* and *K*(*p*) = *K*(*q*) = 0, then *a*(*p, q*) = 1. Thus we add the following constraint:

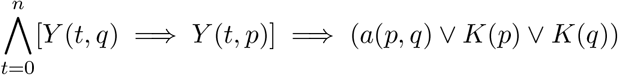 In order to implement this efficiently in Conjunctive Normal Form (CNF) required by all available wMax-SAT solvers, we need to introduce additional boolean variables *a_t_*(*p, q*) for all values of *t* = 0, . . . , *n*. For each row *t* (including 0) and each pair of columns *p* and *q* (including *x*) we must thus satisfy the constraint:

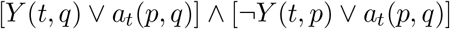

and the final constraint that must be satisfied for each pair *p, q* is:

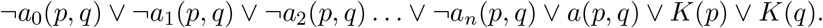
3. For all triplet of mutations *p, q* and *r* such that *p* is an ancestor of *q* and *r*, but *q* and *r* do not have an ancestor descendant relationship (i.e. they belong to different lineages in the tree), we must satisfy vaf(*p*) · (1 + *δ*) ≥ vaf(*q*) + vaf(*r*). In order to express this as a boolean constraint we introduce yet another boolean function Tvaf(*p, q,r*) defined for all triplet of mutations *p, q, r* (as a part of the input specification) as follows:

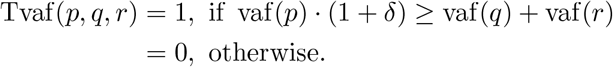 Then the constraint that must be satisfied for all mutations *p, q, r* is:

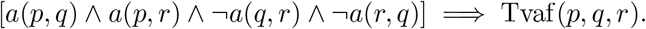

## 3 Results on Simulated Data

We generated simulated data sets and benchmarked our models against alternative tools using three distinct measures of accuracy as described below. We also implemented PhISCS-B using several weighted Max-SAT non-commercial solvers and compared their running time performance with the best performing solver for PhISCS-I showing a significant advantage of using the former. These results suggest that CSP approach can be competitive time-efficient alternative strategy for solving some of the existing problems in tumor phylogenetics which are expressed in the form of integer-linear programs and typically solved by some of the available commercial ILP solvers.

A detailed description of generating simulated data and running the tools is provided in the Appendix.

### 3.1 Comparative Running Time Analysis of PhISCS-B and PhISCS-I

There are a number of available constraint satisfaction software that could be used for our purposes. In order to identify the best performing CSP solvers, we evaluated the top-performing tools from the 2017 Max-SAT competition^3^ (a well-known competition that has been running for many years) on simulated data for the limited version of the problem (with no ISA violations). The competition has both unweighted and weighted tracks. The three top-performing Max-SAT solvers in the weighted competition were MaxHS, QMaxSAT and Maxino. In addition, two other available tools, CPLEX/ILOG by IBM Research and Z3 by Microsoft Research have been benchmarked by the competition organizers. Among these tools, CPLEX/ILOG consistently performed the worst in Max-SAT; this was our experience as well. As CPLEX/ILOG is also commercial and due to its poor performance, here we do not present results obtained by using this tool. Comparisons of running times of PhISCS-B implementations using each of Z3, MaxHS and Maxino Max-SAT solvers are shown in Table 1. This table also contains results of the top-performing ILP (Gurobi) solver that was used to run PhISCS-I.

**Table 1:** Comparison of running times (in seconds) of PhISCS implementations by the use of top-performing CSP solvers from the Max-SAT competition. Results of the top-performing ILP (Gurobi) solver that was used to run PhISCS-I are also included. Runs were performed using the formulation which considers SCS data as the input under infinite sites model. All results are obtained on a single core with a time limit of 24 hours. The number of instances solved within this limit is also reported. Average running times over terminated instances are shown, rounded to the nearest integer, except for the cases where the average is lower than 10 seconds. SC: number of subclones, FN: false negative rate.

| SC | Solver | FN=0.05 |  | FN=0.10 |  | FN=0.15 |  | FN=0.25 |  |
| --- | --- | --- | --- | --- | --- | --- | --- | --- | --- |
|  |  | # inst.<br>solved | Avg.<br>time (s) | # inst.<br>solved | Avg.<br>time (s) | # inst.<br>solved | Avg.<br>time (s) | # inst.<br>solved | Avg.<br>time (s) |
| 4 | ILP (Gurobi) | 10 | 413 | 10 | 1551 | 5 | 1869 | 2 | 5713 |
|  | Z3 | 10 | 29 | 10 | 59 | 8 | 101 | 5 | 347 |
|  | MaxHS | 10 | 39 | 8 | 74 | 9 | 472 | 4 | 5640 |
|  | Maxino | 10 | 4.6 | 10 | 19 | 8 | 1417 | 6 | 26795 |
| 7 | ILP (Gurobi) | 10 | 98 | 10 | 163 | 10 | 110 | 10 | 173 |
|  | Z3 | 10 | 19 | 10 | 28 | 10 | 21 | 10 | 36 |
|  | MaxHS | 10 | 20 | 10 | 29 | 10 | 29 | 10 | 57 |
|  | Maxino | 10 | 2.8 | 10 | 3.8 | 10 | 3.4 | 10 | 6.5 |
| 10 | ILP (Gurobi) | 10 | 56 | 10 | 95 | 10 | 57 | 10 | 71 |
|  | Z3 | 10 | 19 | 10 | 28 | 10 | 24 | 10 | 39 |
|  | MaxHS | 10 | 19 | 10 | 27 | 10 | 25 | 10 | 49 |
|  | Maxino | 10 | 2.7 | 10 | 3.3 | 10 | 2.2 | 10 | 3.9 |

Our results in Table 1 show that, in terms of running time, PhISCS-B significantly outperforms PhISCS-I in all cases. Among the used Max-SAT solvers, Maxino was the top-performing one typically terminating in a few seconds. The only exceptions are cases with 4 subclones and higher false negative error rates. However, even in these computationally most difficult cases, it terminates (within a given time limit) on more instances than most of the other tools which in part explains its higher average running time compared to the other tools (as the running time average was taken only over the terminated instances). While the average running time of Z3 and MaxHS is higher in comparison to Maxino, these tools also show significantly better performance than the Gurobi implementation of PhISCS-I.

### 3.2 Measures of the inference accuracy

In order to assess the (dis)similarity between *G*, the simulated ground truth tree, and *T*, the tree inferred by any one of the methods, we use three measures with distinct properties.

1. **Number of eliminated (ISA violating) mutations**: both true positives (TP) and false positives (FP) in the inferred tree in comparison to the ground truth are considered.
2. **Multi-labeled tree edit distance (MLTED)**: defined as the minimum number of label deletions, empty leaf deletions and vertex expansions applied in any order to transform each of two trees to a maximal common tree. For the practical reasons, here we have used MLTED similarity measure which trivially follows from MLTED. For more details about MLTED and this similarity measure we refer to [33].
3. **Heuristic based Multi-labeled tree edit distance (HMLTED)**: The General Tree Edit Distance TED(*τ, ξ*) between two rooted, unordered, and node-labeled (with a single label) trees *τ* and *ξ* is defined as the minimum number of *edit operations* (insertion, deletion and substitution of nodes and thus their labels) to transform one tree to the other [34, 35]. Here we generalize this notion to *multi-labeled trees,* where each node may have more than one label but each label is unique to a particular node. On a pair of such trees T and G, we introduce Heuristic Multi-Label Tree Edit Distance as HMLTED(*T, G*) = TED(*T*′, *G*′) where *T*′ = *F*(*T*), *G*′ = *F*(*G*) and *F* = argmin_*f*_ TED(*f*(*T*), *f*(*G*)) among all functions *f* which are injective transformations from multi-labeled trees to singly-labeled trees in a way that each multi-labeled node with *k* labels is replaced by a path of uniquely labeled nodes of length *k* (necessarily such transformations will satisfy TED(*f* (*T*), *f* (*G*)) ≥ HMLTED(*T, G*)). Note that we focus on trees *T* and *G* that have the same set of labels. Without loss of generality, we can assume that no pair of labels *x, y* are simultaneously present in a single node in *T* and in *G*, since *x* and *y* can be iteratively replaced with a single new label *v* in both trees. We can also assume that both *T* and *G* have the same root node with identical labeling unique to the root. Since the true tree *G* is typically more compact than inferred tree *T*, we define the transformation *f* first on *G* and then on *T: f* transforms the multi-labeled tree *G* to a singly-labeled tree *G*′ based on *T* in the following way. Consider a breadth first traversal of *G*. Suppose that node *n* visited at a given iteration *i* has labels *L* = {*l*_1_, *l*_2_,…, *l_x_*} in *G*. Now consider all nodes in *T*, namely {*m*_1_, *m*_2_,…, *m_x_*}, whose labels collectively form *L*′ = {*l*_1_, *l*_2_,…, *l_y_*}, the smallest superset of *L*. Then *f* will split *n* into a path of exactly *x* singly (and uniquely) labeled nodes *n*_1_, *n*_2_,…, *n_x_*, with corresponding labels *l*_1_, *l*_2_,…, *l_x_* in the order these labels appear in the breadth first traversal of *T*. Given the transformation *f* from *G* to *G*′ based on *T*, the transformation *f* from *T* to *T*′ based on *G*′ is defined in a similar manner. Interestingly, we prove that this transformation *f* = *F*, i.e. HMLTED(*T, G*) = TED(*f*(*T*), *f*(*G*)) (Theorem 1 in Appendix B).

Note that mutations violating ISA are excluded from these measures.

### 3.3 Accuracy Analysis for PhISCS and Available Methods

We compared both the ILP and CSP implementations of PhISCS against two of the methods operating on single-cell data, namely SCITE and SiFit. We were not able to compare against OncoNEM since it terminated with an error for most of the input matrices nor against ddClone which does not infer phylogeny.

As expected, PhISCS-B and PhISCS-I produce highly similar results with the same value of the objective in all cases and slight differences in the resulting genotypes corrected matrix (denoted as *Y* in Section 2.1). These slight differences are very likely consequence of the existence of multiple optimal solutions in some of the cases. All results presented in this section are obtained by taking the average over 10 simulations we generated for each combination of parameters (see Appendix A for details of simulations).

In Figure 1 we present results for the case where ISA violations are allowed, but no bulk data is used. We focus only on the mutations violating ISA and provide the number of True Positive (TP) and False Positive (FP) calls for such mutations. As our results show, PhISCS outperforms SiFit, the only available alternative method operating under the finite sites assumption, especially in TP measure.

**Figure 1:**
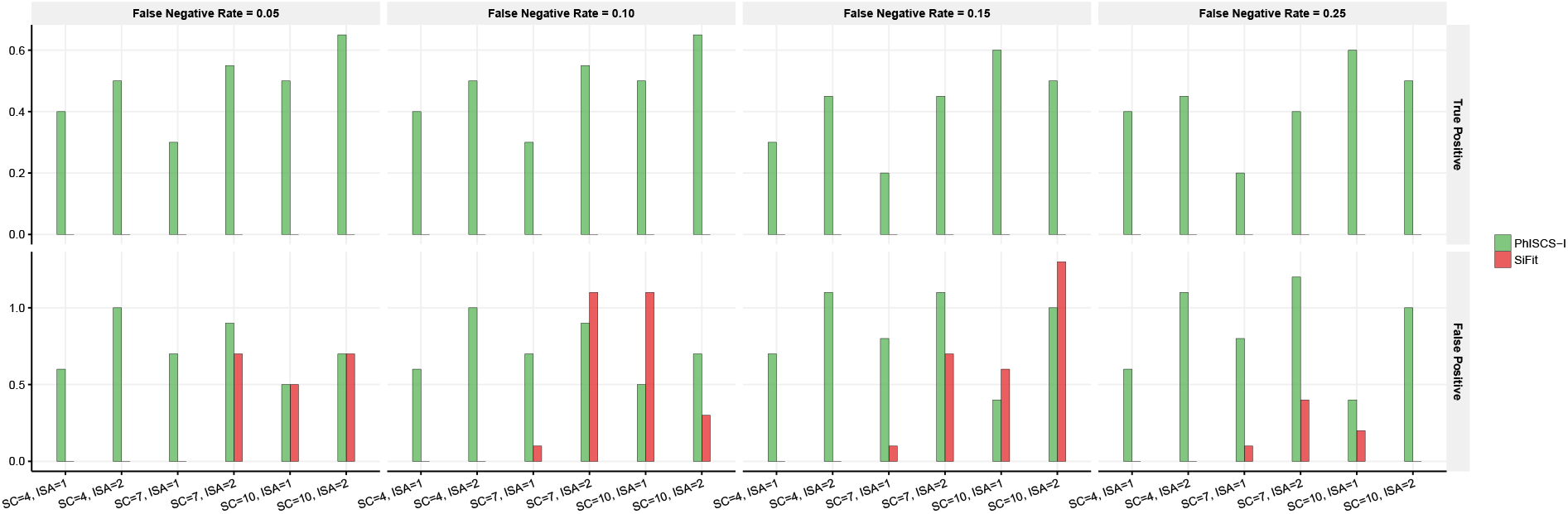
Simulation results for the case where ISA violations are allowed, but only SCS data used as the input. The number of correctly identified ISA violating mutations (True Positive, abbreviated as TP), as well as the number of mutations incorrectly reported to violate ISA (False Negative, abbreviated as FP) in comparison to the ground truth are presented. SC: number of subclones, ISA: number of ISA violating mutations.

Results of PhISCS for the case where both SCS and bulk data are used under finite sites model are presented in Figure 2. (Note that neither SCITE, nor SiFit exploits VAFs obtained from the bulk data.) As Figure 2 illustrates, the combined use of single-cell and bulk data results in improved accuracy for both, TP and FN calls.

**Figure 2:**
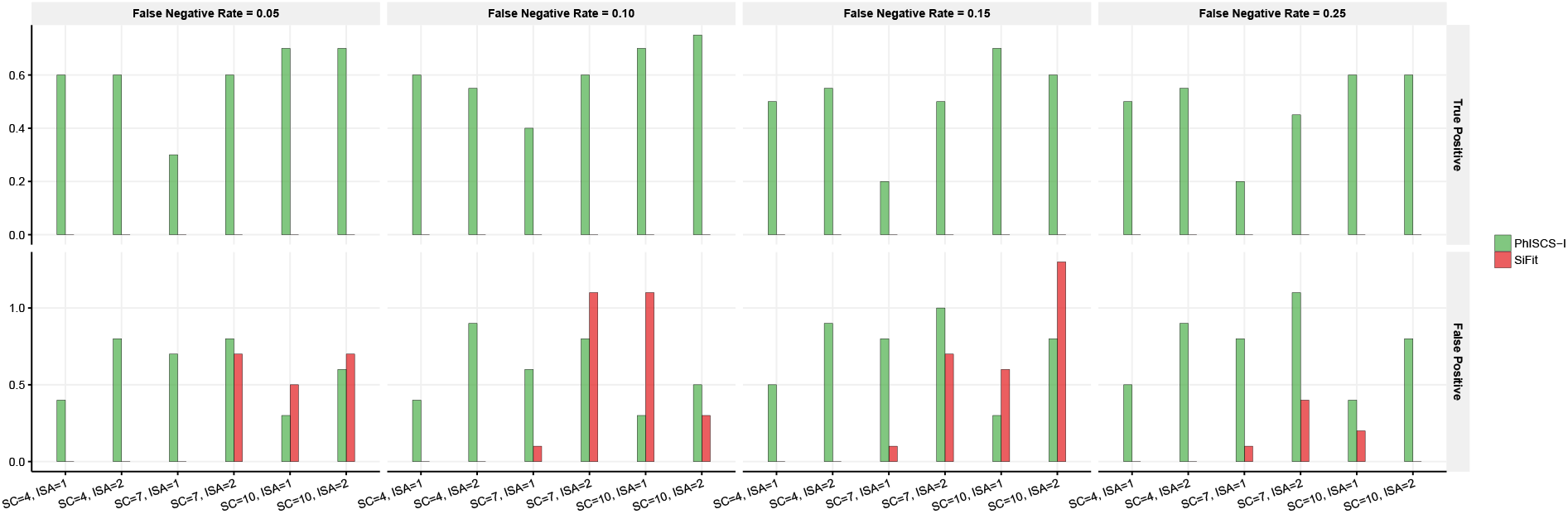
Simulation results ISA violations are allowed and both, single-cell and bulk data used as the input. The number of correctly identified ISA violating mutations (True Positive, abbreviated as TP), as well as the number of mutations incorrectly reported to violate ISA (False Negative, abbreviated as FP) in comparison to the ground truth are presented. SC: number of subclones, ISA: number of ISA violating mutations.

We present the results of PhISCS-I, PhISCS-B and SCITE based on MLTED in Figure 3. In order to facilitate the interpretation of results, we use MLTED similarity measure instead of directly presenting MLTED values. In Figure 3a, we show the results of PhISCS-I and PhISCS-B when only single-cell data is used as the input under infinite sites model. We observe that all three methods: PhISCS-I, PhISCS-B and SCITE have comparable accuracy in this case. Next, we show results of PhISCS when ISA violations are allowed, but no bulk data is used. We can observe that the results of PhISCS-I improve in comparison to the previous case, and overall it has slightly better average performance compared to SCITE. When both bulk and SCS data are provided as the input to PhISCS, we observe further improvements in the accuracy and in this case PhISCS clearly dominates SCITE in terms of the similarity values. Due to the nearly identical performance of PhISCS-B and PhISCS-I we include only results of PhISCS-I in Figure 3b and Figure 3c. We do not include results of SiFit in any of these plots due to its very poor performance in each of the cases.

**Figure 3:**
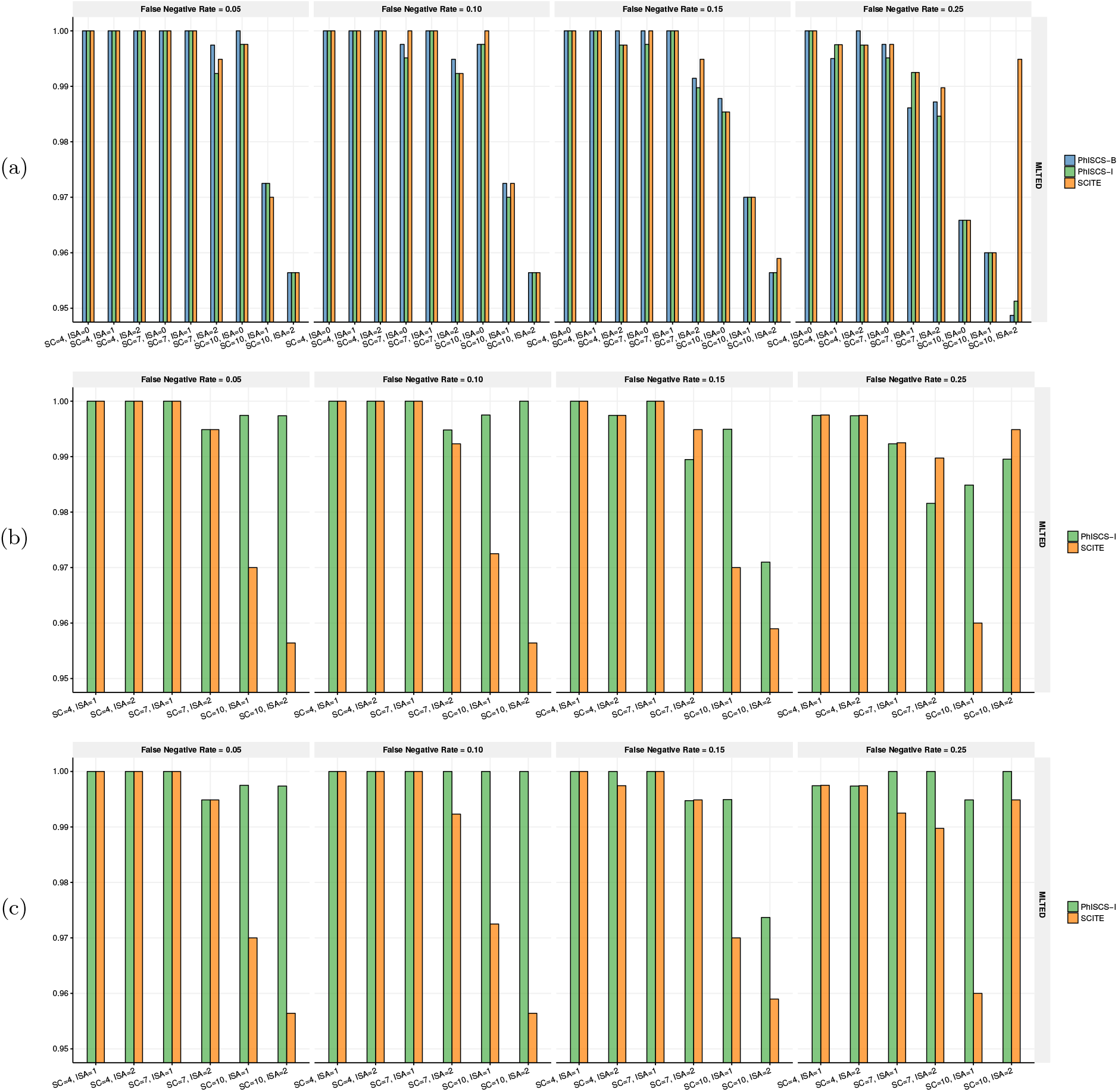
Comparison of different methods based on MLTED similarity measure is shown where values are normalized between 0 to 1. (a) MLTED similarity measure results when no ISA violations are allowed and no bulk data used. (b) MLTED similarity measure results when ISA violations are allowed in PhISCS, but bulk data is not part of the input. (c) MLTED similarity measure results when PhISCS employs both ISA violations and VAFs. Due to the nearly identical performance of PhISCS-B and PhISCS-I in each of (b) and (c) we show only results of PhISCS-I. Due to the very poor performance of SiFit, we do not include any of the results reported by this tool. These plots illustrate that PhISCS-B and PhISCS-I have comparable performance to SCITE in the cases where SCS data is the only input used, however addition of bulk data to the input results in improved performance of PhISCS which clearly outperforms SCITE.

Finally, we discuss results of all tools on our newly introduced HMLTED measure. In Figure 4a we present results of PhISCS-B, PhISCS-I and SCITE for the case where SCS data is used as the only input and infinite sites assumption is made for all mutations (i.e. no mutation elimination is allowed). Results for the runs where ISA violations are allowed are shown in Figure 4b, where only SCS data was used as the input, and in Figure 4c where PhISCS-B was provided with both single-cell and bulk data. As our results illustrate, both implementations of PhISCS and SCITE have comparable accuracy for the case where single-cell data is used as the only input (we suspect that most of the differences are due to non-convergence of MCMC chain in SCITE or multiple equally likely solutions for PhISCS-B and PhISCS-I), however PhISCS significantly benefits from the use of bulk data showing same or improved performance over SCITE in all of the cases under finite sites model. Due to its very poor performance in each of the three cases, we did not include results of SiFit in any of these plots. Also, due to the high similarity of results of PhISCS-B and PhISCS-I, in Figure 4b and Figure 4c we only show results of PhISCS-I.

**Figure 4:**
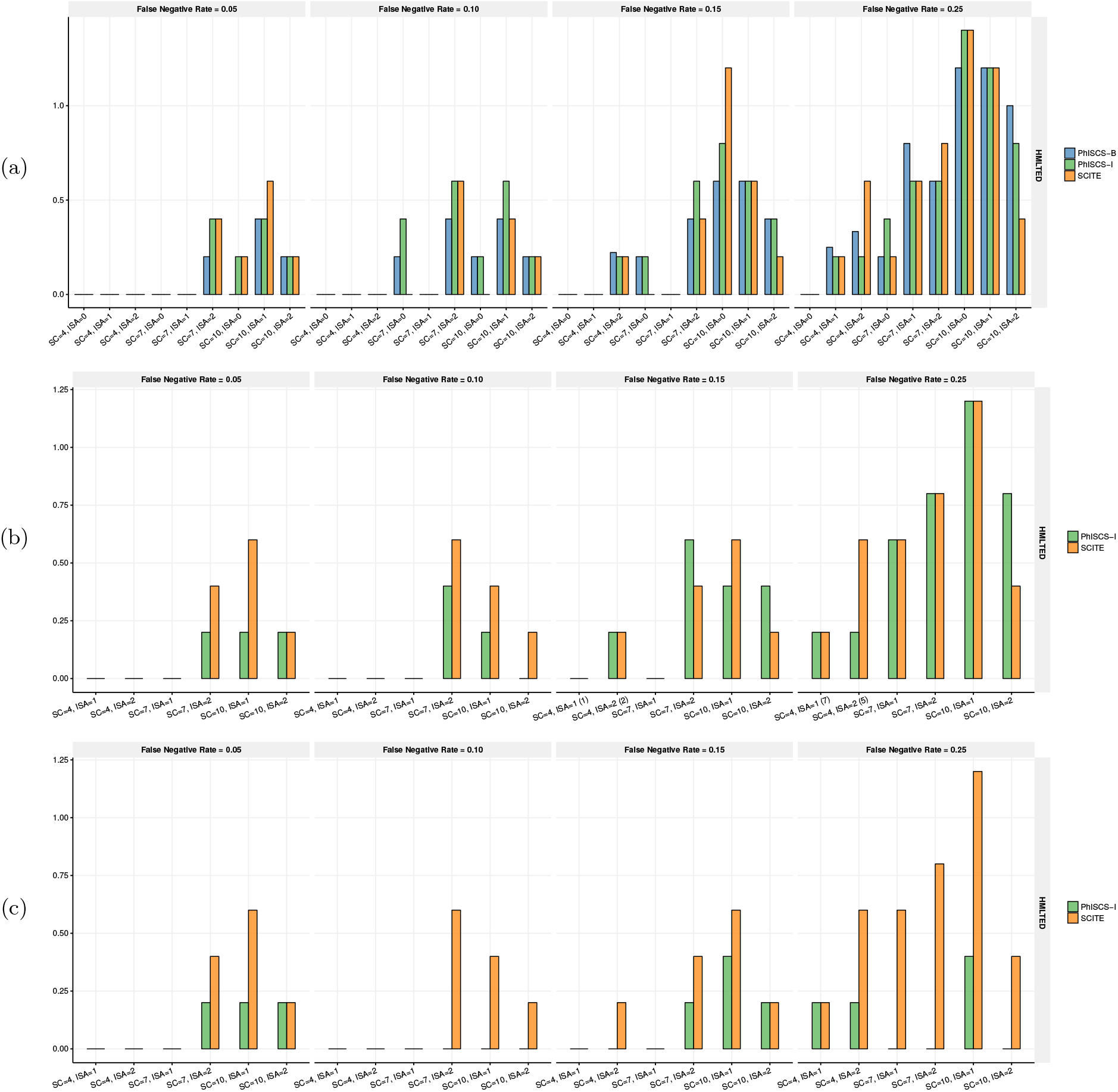
Comparison of different methods based on HMLTED dissimilarity measure. (a) HMLTED values when no ISA violations are allowed. (b) HMLTED values when ISA violations are allowed in PhISCS, but SCS data is used as the only input. (c) HMLTED values when ISA violations are allowed in PhISCS and both single-cell and bulk data are used as the input. CSP and ILP implementations of PhISCS show highly similar performance under finite sites model and therefore in each of (b) and (c) we opted to present only results of PhISCS-I. As our results illustrate, PhISCS-B and PhISCS-I have comparable performance to SCITE in the cases where SCS data is the only input used, however addition of bulk data to the input results in improved performance of PhISCS which outperforms results of SCITE.

## 4 Results on Real Sequencing Data

To demonstrate the utility of our models, we applied PhISCS to two real SCS datasets from recent studies. One of these data sets also provides additional bulk sequencing data with VAF values.

### JAK2-negative myeloproliferative neoplasm

We first run PhISCS on JAK2-negative myeloproliferative neoplasm (essential thrombocythemia) dataset which contains only SCS data [36]. The original study identified 712 SNVs in 58 cells, with an estimated allelic dropout rate of 43.03% and a false discovery rate of 6.04 × 10^-5^. A subset of 18 out of 712 SNVs was identified to be important to tumor progression and retained for further analysis. We used PhISCS to infer tumor phylogeny of the reduced-size dataset which contains 45% of missing values. PhISCS identifies three mutations that break ISA and its output tree topology is provided in Figure 5. Note that this phylogeny does not fully agree with that obtained by earlier studies. Partially this could be due to the lack of VAF values that PhISCS makes extensive use of for the purpose of refining the phylogeny. Furthermore, this data set is very noisy with many missing entries.

**Figure 5:**
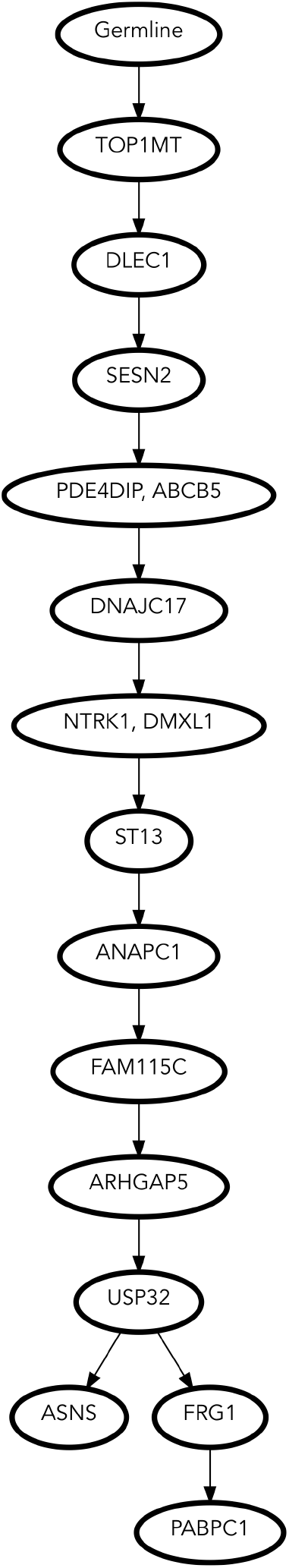
Tree inferred by PhISCS for JAK2-negative myeloproliferative neoplasm. As expected, PhISCS produces a near-linear topology for the tumor phylogeny.

### Childhood acute lymphoblastic leukemia

A more interesting dataset that we tested PhISCS on is obtained from a lymphoblastic leukemia study where both single-cell and bulk sequencing data were made available [37]. We focused on the second patient from this study which has multiple subclones with a non-linear tree topology. For this patient, 16 mutations in 115 single cells were identified. The estimated FN rate is 0.181749. Since SCS data presented in the study are affected by the presence of *doublets,* we first pre-processed SCS data matrix by the use of *Single Cell Genotyper* [14] . The tree inferred by PhISCS, interestingly by the use of SCS data only and with no tolerance for ISA violation (results are shown in Figure 6) seem highly accurate and in good agreement with the tree topology published in the study. The most notable difference is mutation *LINC00052* was placed as a single mutation at branch descending from a clonal population. However, this placement is strongly supported by mutual mutation presence in single-cells where this mutation is rarely present together with some other mutations from the dataset, except for clonal mutations *PLEC, RIMS* and *SIGLEC* as is expected according to the inferred topology. It is interesting to observe that the VAFs of specific mutations (even though they have not been used by PhISCS) seem to contradict with the tree topology (e.g. the mutation on CMTM8). This is likely due to the poor VAF estimation - likely due to an undetected CNV involving the gene.

**Figure 6:**
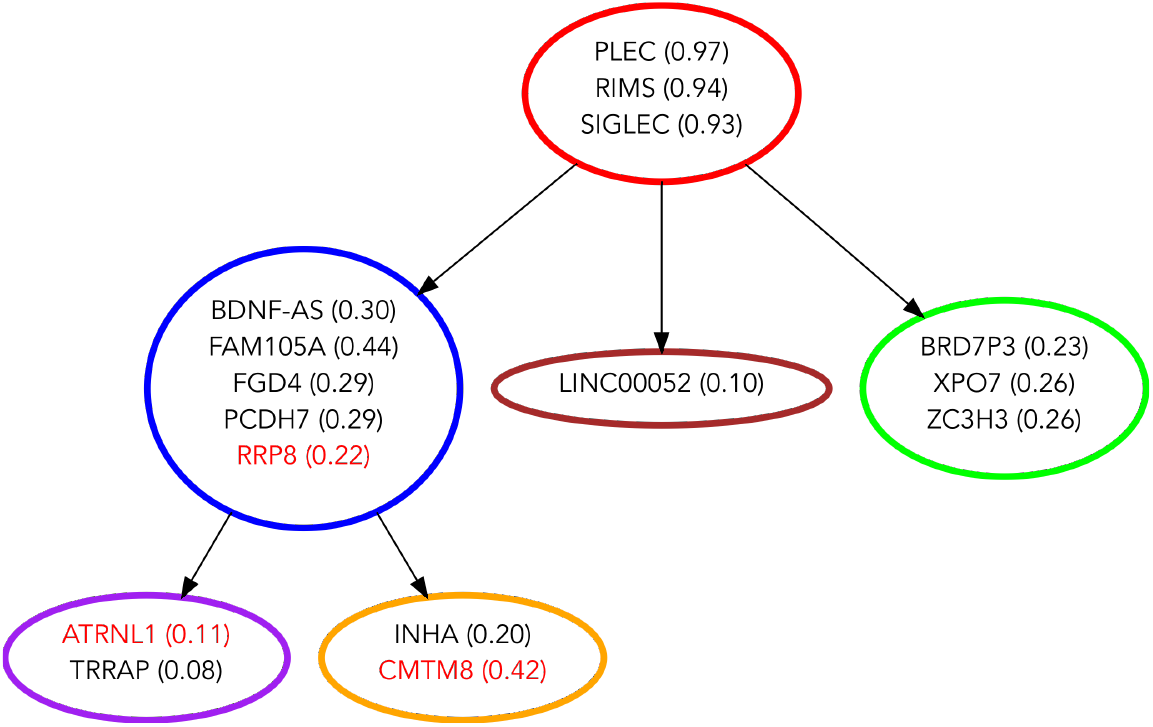
Inferred tumor phylogeny for Patient 2 from [37] through the use of single-cell data only. VAFs derived from bulk data for each mutation (even though they are not used by PhISCS) is provided next to each gene label.

Note that even when we use VAFs and allow ISA violations, we obtain a similar topology as depicted in Figure 7. For this topology, the VAF values do not present a contradiction – which is a highly desirable feature when the VAF values are measured reasonably accurately. Also, note that this particular solution has eliminated three mutations including those in CMTM8 and RRP8. CMTM8 has the mutation with a high VAF value that contradicted with its lower subclonal placement in Figure 6. PhISCS has correctly eliminated it to avoid this contradiction. RRP8, on the other hand, has a mutation whose VAF is much lower than the sum of those in its two child nodes. PhISCS is again correct in eliminating it and thus avoiding any contradiction. All of this points out that PhISCS can potentially help detect previously underexplored copy number aberrations or identify inconsistencies in CNV calls.

**Figure 7:**
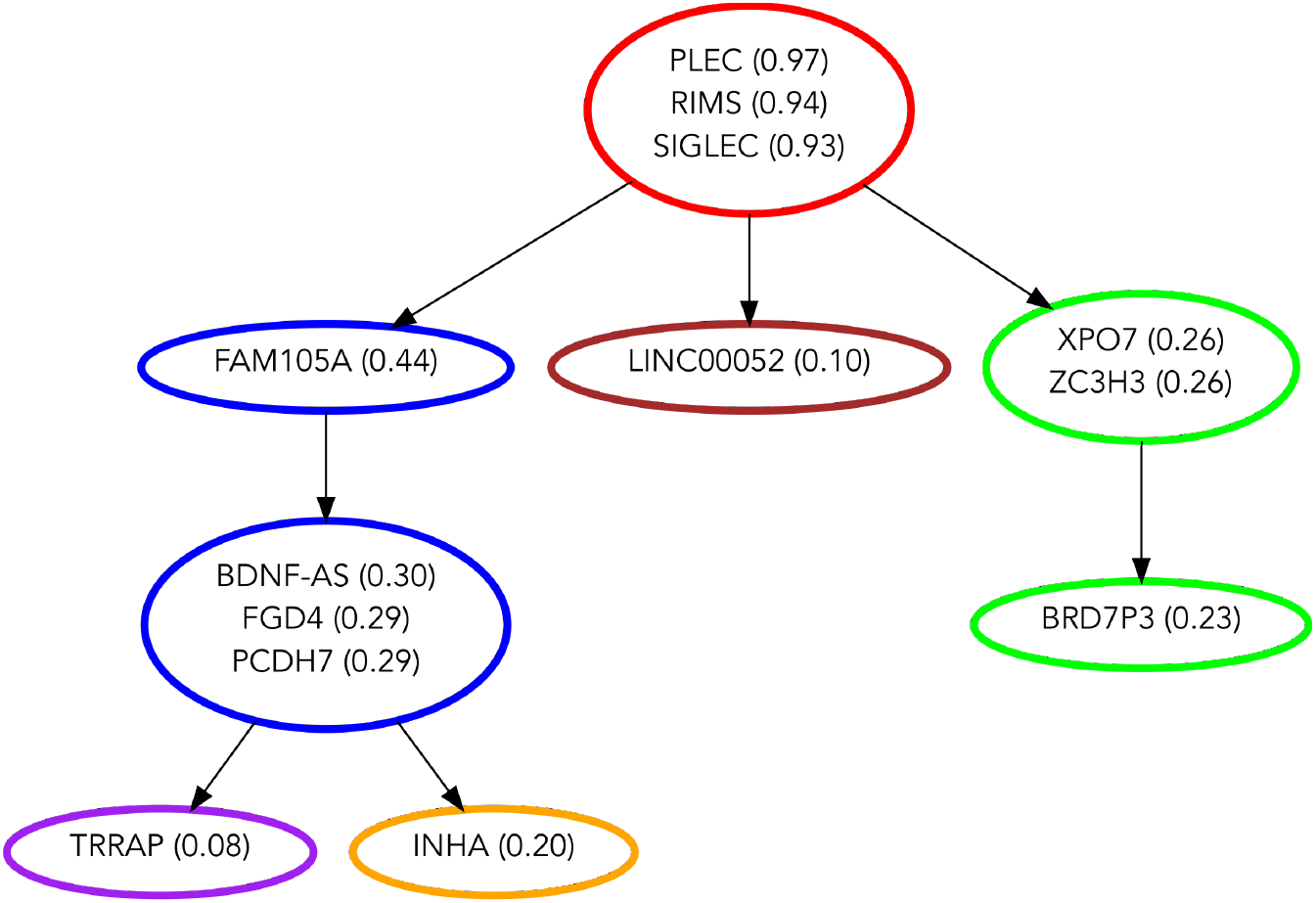
Inferred tumor phylogeny for Patient 2 from [37] through the joint use of bulk and SCS data and allowing ISA violations. The topology of the tree and the mutational placements are highly similar to that in Figure 6 with the exception of three mutations eliminated, possibly due to the inconsistent cellular prevalence and VAF values. Coloring of subclones (nodes) is guided by Figure 6. Subclones corresponding to blue and green nodes are each split into two subclones compared to Figure 6. This is due to the use of VAFs that provide more refined tumor subclonal composition and separation of subclonal populations, otherwise not distinguishable by the use of SCS data only.

## Acknowledgements

SCS and IH would like to acknowledge the Algorithmic Challenges in Genomics program at Simons Institute in UC Berkeley which facilitated this collaboration. IH acknowledges David C. Danko for fruitful discussions on the formulations. This work was supported in part by Indiana University Grand Challenges Program Precision Health Initiative, NSERC Discovery Frontiers Grant, the Cancer Genome Collaboratory to SCS and a Vanier Canada Graduate Scholarship to SM. SC acknowledges a Mobility Exchange Fellowship, which allowed him to visit IH’s group at Weill Cornell. CR and DS were also supported by the Tri-Institutional Training Program in Computational Biology and Medicine (via NIH training grant 1T32GM083937). This work was also supported by start up funds (Weill Cornell Medicine) to IH.

## A Simulation Models used for Benchmarking Tumor Phylogeny Inference Methods

We can depict the history of tumor progression and subclonal composition by the following (the notation is borrowed from [19]):

- Rooted phylogenetic tree *T* with the set of nodes *S*(*T*) of size *s*, where the root node represents the population of healthy cells and each of the remaining nodes represents a distinct population of tumor cells (i.e. a subclone) emerging through selective sweeps during the course of tumor evolution. This tree can be represented by the ancestry matrix *A_T_* defined as follows:

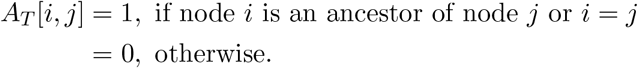
- Set *M* = {*M*_1_, *M*_2_, …, *M_m_*} representing mutations that occur during the course of tumor evolution.
- Function *N_T_* : *M* → *S(T)* where *N_T_*(*x*) denotes the node (i.e. subclone) in tree *T* where mutation *x* occurs for the first time.
- Set *F* = {*f*_1_, *f*_2_,…, *f_s_*} where, for each *i* ∈ {1,2,…, *s*}, *f_i_* is a non-negative real number that represents the frequency of the cellular population corresponding to node *i* of tree *T*. Obviously, frequencies *f_i_* must satisfy 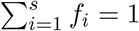.

In our simulations, we restrict ourselves to heterozygous single-nucleotide variants (SNVs) from a diploid regions of the genome.

Under the ISA, for a mutation *M_i_* we define its cellular prevalence *h*(*M_i_*) as a sum of frequencies of cellular populations harboring *M_i_*. According to our notation, *h*(*M_i_*) can be expressed as

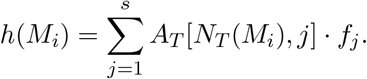

Finally, for node *t* ∈ *T* we define its genotype, denoted as *G_t_*, as a row binary vector of length m such that *G_t_*[*i*] = 1 only if *M_i_* is present in the subclone corresponding to node *t*, which is equivalent to mutation *M_i_* emerging at the node on the path between the root and node *t* (inclusively). More formally, *G_t_*[*i*] = *A_T_*[*N_T_*(*M_i_*),*t*].

Based on the above notation, and using model of clonal tumor evolution and bulk-data simulation presented in [9] and [19], we generate tree *T* of size *s* by randomly choosing one of the possible rooted tree topologies of size s (that were made available in [9]). We then choose values of *f_i_* by formula

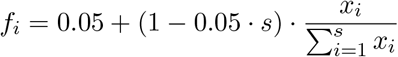

where *x_i_* for *i* ∈ {1, 2,…, *s*} are randomly chosen real numbers from the interval (0,1). The above constant 0.05 ensures the minimal subclonal frequency of 5%. Finally, we randomly spread 40 mutations across the nodes of T, excluding root, and such that each node gets assigned at least one mutation in order to avoid nodes with identical genotypes. We repeat the above simulations 10 times for each *s* ∈ {4, 7,10}.

For mutation *M_i_* we simulate bulk-sequencing read counts from binomial distribution with parameters 5000 (number of trials) and 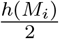 (success probability), where 5000 represents sequencing coverage. Value of *h*(*M_i_*) is divided by 2 in the success probability parameter of binomial distribution due to the assumption that *M_i_* is heterozygous SNV from diploid region of the genome.

For a given tree of tumor evolution and subclonal frequencies simulated above, SCS data is simulated by first drawing 100 single cells from *s* – 1 cancerous populations proportional to their subclonal frequencies and using subclone mutational profiles encoded by row vectors *G_t_*, where *t* ∈ *S*(*T*). After that, we first simulate missing(non-observed) entries of SCS data matrix with rate of 0.10. Next, we simulate false positive entries using the false positive rate of 0.0001. Finally, we introduce false negative sequencing noise by using false negative rates of 0.15 and 0.25 each. This is repeated for all 30 simulations generated above resulting in 60 different simulations.

In order to simulate violations of ISA, for each of the 60 simulations described above, we first generate simulations where exactly one of the mutations violates ISA. In order to do this, we randomly choose mutation *M_i_* and assign it to node *p* different from root and *N_T_*(*M_i_*). Depending on the relation between *p* and *N_T_* (*M_i_*) this violation might represent recurrence or loss of previously obtained mutation. In either case we update genotype of each node and value *h*(*M_i_*) (in an obvious way) and repeat the above procedure of generating bulk and SCS data.

For each of 60 simulations from the previous step, each having exactly one mutation *M_i_* violating ISA, we choose another mutation *M_j_* ≠ *M_i_* from set *M* and apply the same procedure as above simulating two mutations violating ISA.

180 simulations generated in the previous steps are later used as the input for benchmarking.

## B Proof of theorem related to HMLTED measure

### Theorem 1.

*There is a simple transformation f of any input tree T to another T’ and any other input tree I to another I’ for which* TED(*T’, G’*) = HMLTED(*T, G*).

### Proof

We prove this theorem by induction. For the base case, we assume that *T* has two vertices, one being the root and another one vertex with label {*l*_1_} and similarly *G* has two vertices, one being the root and another one vertex with label {*l*_1_}. After applying *f* to *G* based on *T*, we get *G*’ = *G* i.e. *G*’ = *f* (*G*) = *G*. Similarly, after applying *f* to *T* based on *G*’, we get that *T*′ = *T*. So, from the definition of HMLTED, we can write that TED(*T’, G’*) = HMLTED(*T, G*). Thus, the given statement is true.

For the hypothesis step, assume that the given statement is true for *T, G, T’* and *G’*. Now we delete an arbitrary label *I* and corresponding empty vertex from all trees which results in *T*_1_, *G*_1_, 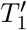 and 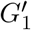, and we assume that 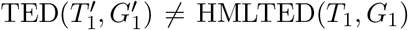. As we apply *f* to *G*_1_ based on *T*_1_ i.e. 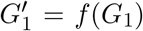 and also apply *f* to *T*_1_ based on 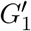, there exists a bijection between the vertices of 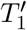 and 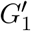. So, a single *l* labelled vertex will be deleted from both 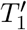 and 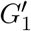. In the transformation of *T*_1_ and *G*_1_ in HMLTED(*T*_1_, *G*_1_), no *l* labelled vertex will exist as *l* is already absent in *T*_1_ and *G*_1_ and it will not change the topology either. Thus there will be no change in distance measure which contradicts our assumption. So, the given statement is true.

## C Details of Running SCITE, SiFit and PhISCS

SCITE and SiFit were both run using default parameter settings, with the exception of chain length in SCITE. In particular, for SCITE we set the number of repetitions to 1 and the chain length of MCMC repetitions to 900000 (10 times higher than default value to allow better convergence). SiFit’s parameters were set to 1 restart and 10000 iterations. Since SiFit outputs cell lineage trees, we had to perform output postprocessing in order to obtain clonal trees of tumor evolution used in our model.

In order to emulate real settings, where only approximate estimates of false negative (FN) and fale positive (FP) error rates are available, in each case we first added noise to true FN and FP error rates and provide resulting noisy values of these parameters as input to each of PhISCS, SCITE and SiFit. The noise was added as follows: if FN error rate used to generate simulations equals β noisy value used as parameter was 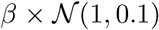, where 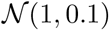 is a random number derived from a normal distribution with mean 1, standard deviation 0.1 and from the interval (0.5, 2) (draws from Normal distribution are repeated in the cases where number falls outside of this range). Analogous was done for adding noise to FP error rates.

In the running simulation for Figure 2 when ISA violations are allowed and both, single-cell and bulk data used as the input, we used *δ* = 0.05.

## D Source codes of Max-SAT solvers used for the implementation of CSP formulation of PhISCS

Source codes of Max-SAT solvers used for the implementation of CSP formulation of PhISCS (i.e. PhISCS-B) are available at:

- Z3: https://github.com/Z3Prover/z3
- MaxHS and Maxino: http://mse17.cs.helsinki.fi/descriptions.html

## Footnotes

1 One additional source of noise is *doublets*, two (or rarely more) cells with heterogeneous mutation profiles treated as a single cell. There already exist computational tools for identifying and decoupling doublets, in particular *Single Cell Genotyper* [14], which we employ in this study as a preprocessing step for the purpose of reducing their impact in our analysis.

2 SAT is very general; it is the very first NP-complete problem.

3 http://mse17.cs.helsinki.fi/index.html

## References

[1] Ash A Alizadeh, Victoria Aranda, Alberto Bardelli, Cedric Blanpain, Christoph Bock, Christine Borowski, Carlos Caldas, Andrea Califano, Michael Doherty, Markus Elsner, et al. Toward understanding and exploiting tumor heterogeneity. Nature Medicine, 21(8):846–853, 2015.

[2] Francesco Strino, Fabio Parisi, Mariann Micsinai, and Yuval Kluger. Trap: a tree approach for fingerprinting subclonal tumor composition. Nucleic acids research, 41(17):e165–e165, 2013.

[3] Wei Jiao, Shankar Vembu, Amit G. Deshwar, Lincoln Stein, and Quaid Morris. Inferring clonal evolution of tumors from single nucleotide somatic mutations. BMC Bioinformatics, 15:35, 2014.

[4] Iman Hajirasouliha, Ahmad Mahmoody, and Benjamin J Raphael. A combinatorial approach for analyzing intra-tumor heterogeneity from high-throughput sequencing data. Bioinformatics, 30(12):i78–i86, 2014.

[5] Amit G Deshwar, Shankar Vembu, Christina K Yung, Gun Ho Jang, Lincoln Stein, and Quaid Morris. Phylowgs: reconstructing subclonal composition and evolution from whole-genome sequencing of tumors. Genome biology, 16(1):35, 2015.

[6] Ke Yuan, Thomas Sakoparnig, Florian Markowetz, and Niko Beerenwinkel. Bitphylogeny: a probabilistic framework for reconstructing intra-tumor phylogenies. Genome biology, 16(1):36, 2015.

[7] Victoria Popic, Raheleh Salari, Iman Hajirasouliha, Dorna Kashef-Haghighi, Robert B West, and Serafim Batzoglou. Fast and scalable inference of multi-sample cancer lineages. Genome Biology, 16(1):91, dec 2015.

[8] Mohammed El-Kebir, Layla Oesper, Hannah Acheson-Field, and Benjamin J Raphael. Reconstruction of clonal trees and tumor composition from multi-sample sequencing data. Bioinformatics, 31(12):i62–i70, 2015.

[9] Salem Malikic, Andrew W McPherson, Nilgun Donmez, and S Cenk Sahinalp. Clonality inference in multiple tumor samples using phylogeny. Bioinformatics, 31(9):1349–1356, 2015.

[10] Nilgun Donmez, Salem Malikic, Alexander W. Wyatt, Martin E. Gleave, Colin Collins, and S Cenk Sahinalp. Clonality inference from single tumor samples using low-coverage sequence data. Journal of Computational Biology, 24(6):515–523, 2017.

[11] Mohammed El-Kebir, Gryte Satas, Layla Oesper, and Benjamin J Raphael. Inferring the mutational history of a tumor using multi-state perfect phylogeny mixtures. Cell systems, 3(1):43–53, 2016.

[12] Francesco Marass, Florent Mouliere, Ke Yuan, Nitzan Rosenfeld, and Florian Markowetz. A phylogenetic latent feature model for clonal deconvolution. Annals of Applied Statistics, 10:2377–2404, 2016.

[13] Gryte Satas and Benjamin J Raphael. Tumor phylogeny inference using tree-constrained importance sampling. Bioinformatics, 33(14):i152–i160, 2017.

[14] Andrew Roth, Andrew McPherson, Emma Laks, Justina Biele, Damian Yap, Adrian Wan, Maia A. Smith, Cydney B. Nielsen, Jessica N. McAlpine, Samuel Aparicio, Alexandre Bouchard-Cote, and Sohrab P. Shah. Clonal genotype and population structure inference from single-cell tumor sequencing. Nat Meth, 13(7):573–576, Jul 2016. Brief Communication.

[15] Katharina Jahn, Jack Kuipers, and Niko Beerenwinkel. Tree inference for single-cell data. Genome Biology, 17(1):86, May 2016.

[16] Edith M. Ross and Florian Markowetz. Onconem: inferring tumor evolution from single-cell sequencing data. Genome Biology, 17(1):69, Apr 2016.

[17] Hamim Zafar, Anthony Tzen, Nicholas Navin, Ken Chen, and Luay Nakhleh. Sifit: inferring tumor trees from single-cell sequencing data under finite-sites models. Genome Biology, 18(1):178, Sep 2017.

[18] Sohrab Salehi, Adi Steif, Andrew Roth, Samuel Aparicio, Alexandre Bouchard-Côté, and Sohrab P. Shah. ddclone: joint statistical inference of clonal populations from single cell and bulk tumour sequencing data. Genome Biology, 18(1):44, Mar 2017.

[19] Salem Malikic, Katharina Jahn, Jack Kuipers, Cenk Sahinalp, and Niko Beerenwinkel. Integrative inference of subclonal tumour evolution from single-cell and bulk sequencing data. bioRxiv, page 234914, 2017.

[20] Dan Gusfield, Yelena Frid, and Dan Brown. Integer Programming Formulations and Computations Solving Phylogenetic and Population Genetic Problems with Missing or Genotypic Data, pages 51–64. Springer Berlin Heidelberg, Berlin, Heidelberg, 2007.

[21] Dan Gusfield. Persistent phylogeny: A galled-tree and integer linear programming approach. In Proceedings of the 6th ACM Conference on Bioinformatics, Computational Biology and Health Informatics, BCB ‘15, pages 443–451, New York, NY, USA, 2015. ACM.

[22] Leslie Ann Goldberg, Paul W. Goldberg, Cynthia A. Phillips, Elizabeth Sweedyk, and Tandy Warnow. Minimizing phylogenetic number to find good evolutionary trees. Discrete Applied Mathematics, 71(1):111 – 136, 1996.

[23] Paola Bonizzoni, Chiara Braghin, Riccardo Dondi, and Gabriella Trucco. The binary perfect phylogeny with persistent characters. Theoretical Computer Science, 454(Supplement C):51 – 63, 2012. Formal and Natural Computing.

[24] Paola Bonizzoni, Simone Ciccolella, Gianluca Della Vedova, and Mauricio Soto. Beyond perfect phylogeny: Multisample phylogeny reconstruction via ilp. In Proceedings of the 8th ACM International Conference on Bioinformatics, Computational Biology,and Health Informatics, ACM-BCB ‘17, pages 1–10, New York, NY, USA, 2017. ACM.

[25] Mario Alviano, Carmine Dodaro, and Francesco Ricca. A maxsat algorithm using cardinality constraints of bounded size. In IJCAI, pages 2677–2683, 2015.

[26] Jessica Davies and Fahiem Bacchus. Solving MAXSAT by solving a sequence of simpler SAT instances. In Principles and Practice of Constraint Programming - CP 2011 - 17th International Conference, CP 2011, Perugia, Italy, September 12–16, 2011. Proceedings, pages 225–239, 2011.

[27] Jessica Davies and Fahiem Bacchus. Exploiting the power of mip solvers in maxsat. In International conference on theory and applications of satisfiability testing, pages 166—181. Springer, 2013.

[28] Jessica Davies and Fahiem Bacchus. Postponing optimization to speed up MAXSAT solving. In Principles and Practice of Constraint Programming - 19th International Conference, CP 2013, Uppsala, Sweden, September 16–20, 2013. Proceedings, pages 247—262, 2013.

[29] Inês Lynce and João Marques-Silva. Efficient haplotype inference with boolean satisfiability. In Proceedings of the AAAI, 2006.

[30] Jost Neigenfind, Gabor Gyetvai, Rico Basekow, Svenja Diehl, Ute Achenbach, Christiane Gebhardt, Joachim Selbig, and Birgit Kersten. Haplotype inference from unphased snp data in heterozygous polyploids based on sat. BMC genomics, 9:356, 2008.

[31] Dan He, Arthur Choi, Knot Pipatsrisawat, Adnan Darwiche, and Eleazar Eskin. Optimal algorithms for haplotype assembly from whole-genome sequence data. Bioinformatics, 26(12):i183–i190, 2010.

[32] Jack Kuipers, Katharina Jahn, Benjamin J Raphael, and Niko Beerenwinkel. Single-cell sequencing data reveal widespread recurrence and loss of mutational hits in the life histories of tumors. Genome research, 2017.

[33] Nikolai Karpov, Salem Malikic, Md. Khaledur Rahman, and S. Cenk Sahinalp. A multilabel tree edit distance for comparing “clonal trees” of tumor progression. To appear in 18th proceedings of Workshop on Algorithms in Bioinformatics (WABI2018). The implementation of this method can be found at https://github.com/khaled-rahman/leafDelTED.

[34] Kaizhong Zhang, Rick Statman, and Dennis Shasha. On the editing distance between unordered labeled trees. Information processing letters, 42(3):133–139, 1992.

[35] Kaizhong Zhang. A new editing based distance between unordered labeled trees. In Combinatorial Pattern Matching, pages 254—265. Springer, 1993.

[36] Yong Hou, Luting Song, Ping Zhu, Bo Zhang, Ye Tao, Xun Xu, Fuqiang Li, Kui Wu, Jie Liang, Di Shao, Hanjie Wu, Xiaofei Ye, Chen Ye, Renhua Wu, Min Jian, Yan Chen, Wei Xie, Ruren Zhang, Lei Chen, Xin Liu, Xiaotian Yao, Hancheng Zheng, Chang Yu, Qibin Li, Zhuolin Gong, Mao Mao, Xu Yang, Lin Yang, Jingxiang Li, Wen Wang, Zuhong Lu, Ning Gu, Goodman Laurie, Lars Bolund, Karsten Kristiansen, Jian Wang, Huanming Yang, Yingrui Li, Xiuqing Zhang, and Jun Wang. Single-cell exome sequencing and monoclonal evolution of a jak2-negative myeloproliferative neoplasm. Cell, 148(5):873–885, 2017.

[37] Charles Gawad, Winston Koh, and Stephen R. Quake. Dissecting the clonal origins of childhood acute lymphoblastic leukemia by single-cell genomics. Proceedings of the National Academy of Sciences, 111(50):17947–17952, 2014.

